# Commonly used non-antibiotic medications promote mutation frequency and antimicrobial resistance in *Escherichia coli*

**DOI:** 10.1101/2025.01.13.632802

**Authors:** Hanbiao Chen, Sylvia A. Sapula, John Turnidge, Henrietta Venter

## Abstract

Antimicrobial resistance (AMR) poses a global threat to public health. The excessive use of antibiotics significantly contributed to the rise of resistance. Recent evidence suggests that non-antibiotic medications (NAMs) also play a role in antimicrobial resistance development, although this aspect remains less explored and understood. This issue is particularly relevant in residential aged care facilities (RACFs) where both NAMs and antibiotics are frequently used, and AMR is prevalent. We investigated the propensity of NAMs that are commonly used in RACFs and contribute to polypharmacy including non-steroidal anti-inflammatory drugs (NSAIDs) such as ibuprofen and diclofenac, acetaminophen (antipyretic), metformin (glucose-lowering medication), atorvastatin (lipid-lowering agent), tramadol (opioid analgesic), temazepam (hypnotic), and pseudoephedrine (sympathomimetic) to promote bacterial antibiotic resistance by increasing the acquisition of mutations. *Escherichia coli* was exposed to different NAMs at their gut concentration, combined with ciprofloxacin and the mutation frequency was determined. Additionally, we explored the simultaneous effect of two NAMs as a starting point for studying polypharmacy.

Ibuprofen and acetaminophen significantly increased mutation frequency, and conferred high levels of ciprofloxacin resistance, especially when *E. coli* was exposed to two NAMs. Whole genome sequencing revealed that these changes correlated with mutations in DNA gyrase GyrA, the multiple antibiotic resistance regulator, MarR, and the drug efflux pump expression suppressor, AcrR. Consequently, an increase in transcription of the *acrA* gene from the AcrAB-TolC drug efflux pump was observed. The combination of two NAMs increased the mutation rate. These multiple mutations caused the higher levels of ciprofloxacin resistance that were observed.

Given the risk of polypharmacy to induce AMR, and the results observed in this study, the assessment NAMs in their ability to promote bacterial resistance warrants special attention in future studies into prescribing practices.

## Introduction

The development and spread of antimicrobial resistant organisms and resistance genes represents one of the greatest threats facing public health (WHO, 2015). This is exemplified by current reports estimating that 4.95 million deaths were associated with antimicrobial resistance in 2019 (Murray et al., 2022). It is well-known that the prevalent amplifier behind the development of antimicrobial resistance (AMR) is the overuse of antibiotics in clinical and agriculture settings (Martin et al., 2015; Rousham et al., 2018). However, in addition to antibiotics, the use of non-antibiotic medications (NAMs) such as diuretics, statins, proton-pump inhibitors, amongst others, have also been shown to contribute the rise of AMR (Imhann et al., 2016; Wang et al., 2020).

A recent report assessing antimicrobial properties of NAMs revealed that more than 200 commonly used drugs possessed antibiotic-like effects on gut bacteria (Maier et al., 2018). Currently, the global pharmaceutical market is primarily made up of non-antibiotic drugs which occupy 95% of the market (Hamad, 2010). The pharmaceutical market also continues to grow, experiencing significant growth during the last two years, with worldwide revenues totalling 1.48 trillion (US) dollars in 2022 (Mikulic, 2024). The growth of this industry is not surprising, as the world’s population is ageing (Christensen et al., 2009), and this ageing population relies on an arsenal of medicines to manage chronic disease (Sluggett et al., 2024) with some taking nine or more prescribed drugs per day, a phenomenon known as polypharmacy, and one which is increasing worldwide (Dagli & Sharma, 2014; Sluggett et al., 2024; Wang et al., 2021; Wastesson et al., 2018). Older persons also represent a population with a high consumption of antibiotics, which are frequently overused in residential aged care facilities (RACFs) to manage urinary tract and respiratory infections (ACSQHC, 2024; Fleming et al., 2015; Maher et al., 2024; Nicolle et al., 2000; Sluggett et al., 2021).

NAMs can have both a direct and/or indirect effect on the development of AMR. Some NAMs display direct antibacterial activity, such as statins (Ko et al., 2017). Others, such as certain nonsteroidal anti-inflammatory drugs (NSAIDs), can promote the development of cross-resistance, selecting for a mechanism of resistance which can concurrently confer resistance to antibiotics, best exemplified by multidrug efflux pumps (Verma et al., 2018). Another important effect, which can be considered both direct and indirect, results from the enhancement of transmission of AMR determinants though bacterial transformation, exemplified by NSAIDs such as diclofenac (Wang et al., 2020). Compounding these is the fact that some antibiotics like ciprofloxacin, a fluoroquinolone, and non-antibiotics like tramadol, a weak opioid, are not completely degraded in the body and end up in wastewater through the excretion of urine and/or stool (Bírošová et al., 2020). Once there, these, and the metabolites of both antibiotics and non-antibiotic drugs, exert their combined effect and create a perfect environment for the development and dissemination of AMR.

To assess the propensity of NAMs to promote the development of AMR, we investigated nine medications; acetaminophen, ibuprofen, diclofenac, furosemide, atorvastatin calcium, metformin, pseudoephedrine, temazepam, and tramadol which are commonly used by older people. As these NAMs are frequently co-administered with antibiotics, mutation frequencies were determined in the presence of ciprofloxacin. Ciprofloxacin is not only a known inducer of mutations but is also used in treatment of urinary tract infections (UTIs) which are one of the most common bacterial infections (Blondeau, 2004; Breidenstein et al., 2012). The frequent usage of ciprofloxacin in RACF settings has led to the emergence of ciprofloxacin resistance and ciprofloxacin-resistant *Escherichia coli* which has posed a threat to residents living in RACFs (Fasugba et al., 2015). The effect of the nine NAMs and combinations of two NAMs on the mutation frequencies were determined for two *E. coli* isolates, *E. coli* BW25113, a derivative of *E. coli* K-12 and *E. coli* 6146 that was isolated from a faecal sample of a retirement community resident. The antimicrobial susceptibilities of the NAM-induced mutants and the mechanisms underlying the observed resistance were assessed by whole genome sequencing. The role of drug efflux pumps, which can be over-expressed in response to stress, and contribute to cross-resistance, was also further explored.

## Materials and Methods

### 2.1. Bacterial strains and culture conditions

Bacterial strains used in this study included the *E. coli* BW25113 strain, a derivative of *E. coli* K-12 strain (Datsenko & Wanner, 2000; Grenier et al., 2014), and *E. coli* 6146 originally isolated from a faecal sample obtained from a retirement community in Adelaide, Australia. The project received ethical approval from the University of South Australia’s human research ethics committee to collect faecal samples from RACF and retirement residents (Application ID: 201882). Typically, strains were cultured in Luria-Bertani (LB) media (Difco®, Franklin Lakes, New Jersey) at 37 °C. The NAMs were dissolved in DMSO, and antibiotics were dissolved in their recommended solvents. Nutrient agar (NA) was chosen as the medium for studies, with NA plates supplemented with ciprofloxacin and NAMs to assess mutation frequency.

### 2.2. Culture conditions and growth assays

Growth assays were carried out in a volume of 200 μL of the inoculum in cation-adjusted Mueller Hinton II Broth (BD, Australia). Briefly, cells standardised to OD_600_ of 0.015 were added to each well of a 96-well microtiter plate (Costar®) containing concentration ranges of ciprofloxacin (0, ½, and ¾ the MIC of ciprofloxacin) and each NAM at gut concentration (Maier et al., 2018). The 96 well-plate was incubated for 20 hours at 37 °C in the EnSpire 2300 Multilabel Reader (PerkinElmer, USA). Absorbance at 600 nm was measured and recorded at 20 minutes time intervals.

### 2.3. Estimation of the concentration of NAMs in the intestine

The gut concentration of NAMs was calculated using a model based on Maier et al., (2018). Briefly, this method assumes that an oral drug is dissolved in 300 mL of water including water allowing to swallow the pill (240 mL) and the resting water (60 mL) in the small intestine. The model also assumes that 90% of oral drugs will be absorbed into the bloodstream, meaning that 10% of the drugs will remain in the intestinal area. Single doses will be obtained from the Defined Daily Dose (DDD) database (WHO, 1996).

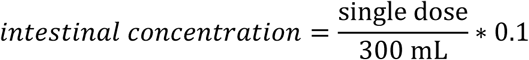

Based on the above model, NAMs concentration used in the study were 216 μg/mL for acetaminophen, 15.4 μg/mL for diclofenac, 13.2 μg/mL for furosemide, 99.8 μg/mL for ibuprofen, 194.6 μg/mL for metformin, 6.6 μg/mL for atorvastatin, 40 μg/mL for pseudoephedrine, 3.3 μg/mL for temazepam, and 33.3 μg/mL for tramadol.

### 2.4. Antimicrobial Susceptibility Testing

Antimicrobials assessed in this study included ciprofloxacin, minocycline, cefepime, meropenem, norfloxacin, levofloxacin, erythromycin, and novobiocin, all obtained from Sigma Aldrich (Sydney, Australia). NAMs assessed included acetaminophen, furosemide, atorvastatin calcium, diclofenac, ibuprofen, and metformin were also used and were also obtained from Sigma Aldrich (Sydney, Australia). Each antibiotic and pharmaceutical compound was dissolved in the appropriate solvent, according to the manufacturer’s recommendations.

Minimum inhibitory concentrations (MICs) were determined using the broth microdilution assay ISO 20776-1 as recommended by European Committee on Antimicrobial Susceptibility Testing (Eucast, 2000). Each strain was tested in triplicate.

### 2.5. Determination of mutation frequency

Mutation frequency was assessed using both the *E. coli* BW25113 (wild type) and *E. coli* 6146 (an antibiotic sensitive isolate from the faecal sample of a retirement resident). Overnight cultures grown at 37 °C in 5 mL LB broth were diluted (1:100) in LB and grown to mid-log phase (OD_600_ of 0.4 - 0.6) at 37 °C. A volume of 100 μL of each culture was spread plated onto NA plates containing 1x, 2x, and 4x the MIC of ciprofloxacin (0.015 to 0.06 μg/mL) and NAMs, at clinically relevant concentrations (Maier et al., 2018), to select ciprofloxacin-resistant bacteria. Cultures were incubated for 48 h at 37 °C. The mutation frequency was calculated according to the following equation:

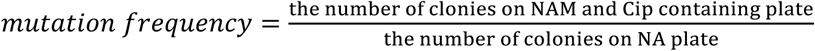

To address plating bias, assays were carried out on four different days with different batches of cells. A representative number (√n+1) of *E. coli* derived mutants were randomly picked and MIC of ciprofloxacin was determined. The representative mutants displaying at least 4-fold increased ciprofloxacin resistance were selected for further analysis zto avoid including strains whose MIC increase may have been due to MIC assay variation.

### 2.6. Checkerboard assays

To determine the effect of RND efflux pump-mediated resistance, checkerboard assays were carried out on ciprofloxacin resistant isolates in the presence and absence of the efflux pump inhibitor Phenylalanine-arginine β-naphthylamide (PAβN) as described previously (Blaikie et al., 2024; Ohene-Agyei et al., 2014; Pisoni et al., 2023). Ciprofloxacin was diluted vertically down the 96-well microtiter plate (Costar®) (2-fold serial dilutions), and PAβN was serially diluted horizontally across the plate (2-fold serial dilutions). Inoculum of 1.5 x 10^6^ CFU/mL was added to each well and plates were incubated at 37 °C for 18 hours, shaking at 120 rpm. The MICs in the presence of PAβN were used to compare the fold change in MICs in the absence of PaβN.

### 2.7. DNA extraction, whole genome sequencing and data processing

Genomic DNA was extracted using the NucleoSpin® Microbial DNA extraction kit (MACHEREY-NAGEL, Germany) according to manufacturer’s instruction. The concentration and purify of DNA (A_260_/A_280_, 1.8–2.0) were assessed using the Cytation 5® (Bio-Tek®, Winooski, USA) plate reader and the quality of the DNA analysed by agarose gel electrophoresis. All samples were then submitted to the Australian Genome Research Facility (AGRF) for library preparation and paired-end sequencing were performed on the Illumina Hi-Seq platform.

Raw 150 base paired-end sequencing reads were assembled and annotated using the TORMES pipeline (v.1.3) (Quijada et al., 2019). To assess possible mutations, draft genomes of *E. coli* derived mutant were screened against the wild-type using BRESEQ (Barrick et al., 2014). Amino acids of target wild type proteins were aligned to mutant amino acid sequences using CLUSTAL O v.1.2.4 (Sievers & Higgins, 2018). Sequence variations were visualized using ESPript v.3.0 (Robert & Gouet, 2014).

### 2.8. RNA extraction, Reverse-Transcription Quantitative Real-Time Polymerase Chain Reaction (RT-qPCR)

Reverse transcription quantitative real-time polymerase chain reaction (RT-qPCR) was carried out to quantify the transcriptional level of *acrA*, which encodes for the AcrA membrane fusion protein of the AcrAB-TolC multidrug efflux pump. Isolates displaying over 4-fold increased ciprofloxacin resistance (MIC of ciprofloxacin ≥ 0.015 μg/mL) were selected for RT-qPCR analysis to assess the potential role of efflux pumps in this resistance.

Overnight cultures inoculated with single colonies were diluted (1:100) in LB broth and grown until mid-log phase (A_600_ of 0.5) at 37 °C. The total RNA was extracted as previously described (Amsalu et al., 2020). Briefly, 1 mL of cultures were centrifugated, resuspended in Trizol and chloroform, and then further purified using the NucleoSpin®RNA extraction kit (MACHEREY-NAGEL, Germany) followed by rDNase digestion which was performed to remove DNA during this period. The concentration and purity of total RNA were quantified through Cytation 5® (Bio-Tek®, Winooski, USA).

All RT-qPCR reactions were performed using a magnetic induction cycler (Bio Molecular Systems, Australia) and the KAPA SYBR® FAST qPCR Master Mix kit (Kapa Biosystems, USA). The housekeeping gene, *recA,* was used to normalise the transcriptional levels of *acrA*, the target gene. The primer sequences include AcrA_F 5′-GATCCAGCCACCTATCAGGC-3′; AcrA_R 5′-ACGCTTGACCGTCAGTTGAT-3′; RecA_F 5′-TTAAACAGGCCGAATTCCAG-3′; RecA_R 5′-CCGCTTTCTCAATCAGCTTC-3′. Cycling parameters were as follows: reverse transcription at 42 °C for 10 min, inactivation at 95 °C for 3 min, and 40 cycles of PCR at 95 °C for 5 s, 60 °C for 30 s, and 72 °C for 15 s. No template control (NTC) and no reverse-transcriptase control (NRT) were established, and the qPCR reaction samples carried out in triplicate.

### 2.9. Statistical analysis

Graph Prism v9. was used for all data analysis. Significant differences were assessed using an independent-sample t-tests and corrected by the Bonferroni correction method, *P < 0.05, **P < 0.01, ***P < 0.001, and ****P < 0.0001. All experiments were conducted in triplicate.

## Results

### 3.1. NAM exposure increases fitness of E. coli in the presence of ciprofloxacin

To assess the propensity of commonly used non-antibiotics to induce bacterial resistance to antibiotics, six NAMs were chosen based on their frequent use among the elderly in RACFs (Juurlink et al., 2003; Kalisch Ellett et al., 2021). Ciprofloxacin was used in this study as it is a known inducer of mutations (Rodriguez-Martinez et al., 2006) with an increase in ciprofloxacin-resistant uropathogens representing a threat to residents living in aged care facilities (Fasugba et al., 2015; Mulder et al., 2016; Sapula et al., 2023; Stuart et al., 2012). Mutation frequencies were determined in *E. coli* BW25113 and in, *E. coli* 6146 (an antibiotic sensitive strain isolated from a retirement resident). The baseline antimicrobial susceptibility of the test strains, *E. coli* BW25113 and the *E. coli* 6146, was determined and shown to be almost identical and wild type for the agents tested (**Table 1**). None of the NAMs assessed in this study displayed antimicrobial activity against these isolates, even at the highest concentration tested (512 μg/mL). To obtain a comprehensive set of antibiotic susceptibility data, erythromycin and novobiocin, to which *E. coli* strains are known to be resistant, were also included (Visvalingam et al., 2017).

**Table 1.**
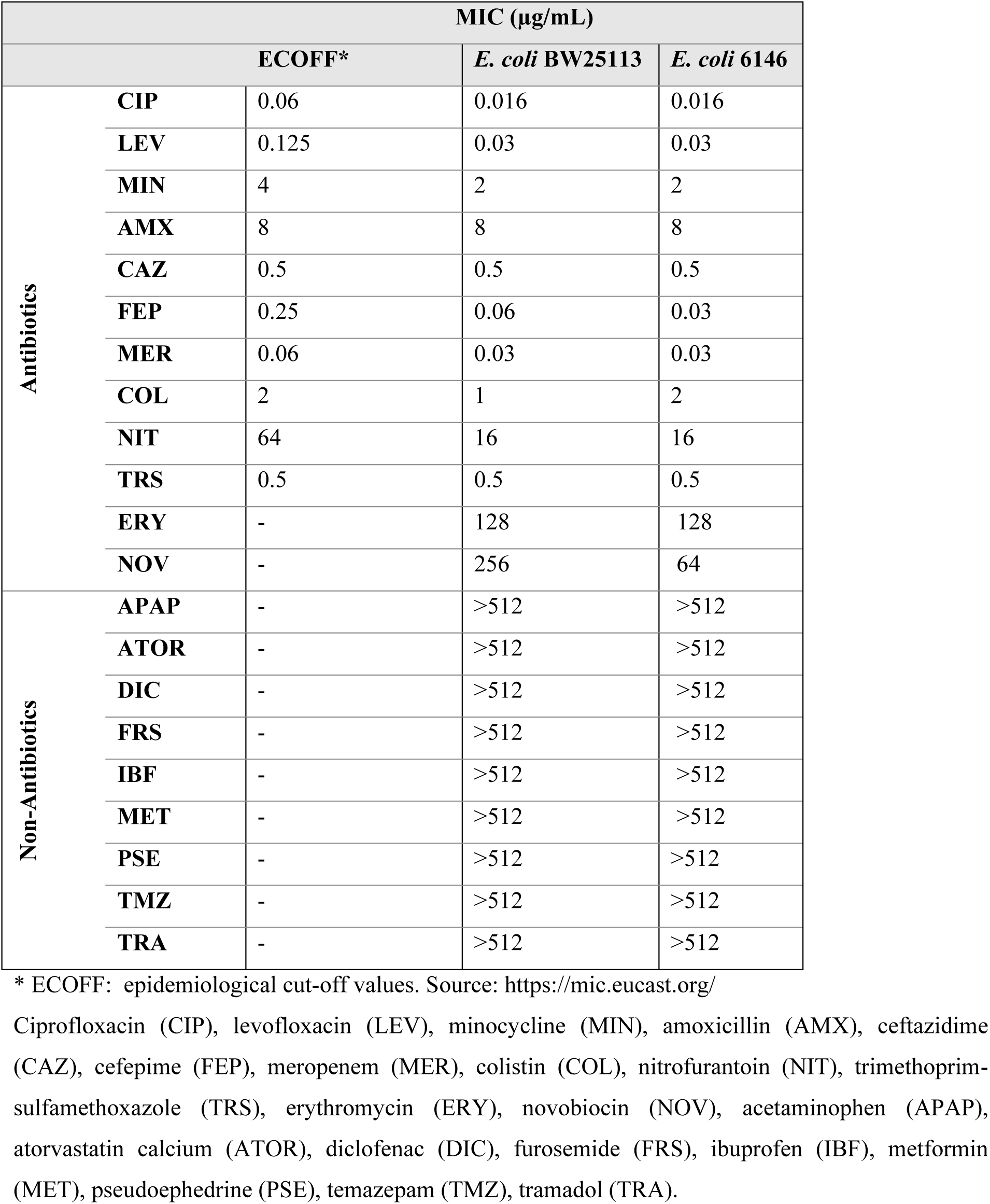
MICs for *E. coli* BW25113 and *E. coli* 6146 against antibiotics and NAMs assessed in this study.

To investigate whether growth was affected by NAM exposure, the growth of *E*. *coli* BW25113 and *E*. *coli* 6146 was analysed in the presence of sub-inhibitory concentrations of ciprofloxacin and/or gut concentrations of the NAMs analysed in this study (**Figure 1**). As expected, when compared to non-exposed cells the addition of NAMs alone did not impact the growth of *E. coli* cells (**Figures 1A and 1B**). However, the growth rate and maximum cell density were dramatically affected at ¾ the MIC of ciprofloxacin for both *E. coli* BW25113 and *E. coli* 6146 cells. Reduction of cell growth was observed from 0.457 ± 0.009 h^-1^ to 0.140 ± 0.006 h^-1^ and 0.443 ± 0.006 h^-1^ to 0.007 ± 0.001 h^-1^ for *E. coli* BW25113 and *E. coli* 6146 respectively (**Figure 1C** and **1D**). Additionally, the start of the exponential growth rate was delayed for *E. coli* 6146 cells in the presence of ciprofloxacin (**Figure 1D**). While in the presence of ¾ the MIC of ciprofloxacin, diclofenac, ibuprofen, and acetaminophen increased the growth rates of *E. coli* BW25113 to 0.201 ± 0.004, 0.179 ± 0.006, and 0.2823 ± 0.013 h^-1^ respectively (**Figure 1C**). The same three NAMs were also able to reduce the length of the lag phase (**Figure 1D**). Collectively, certain NAMs increase the fitness and adaptability of *E. coli* under stress conditions such as growth in the presence of an antibiotic.

**Figure 1:**
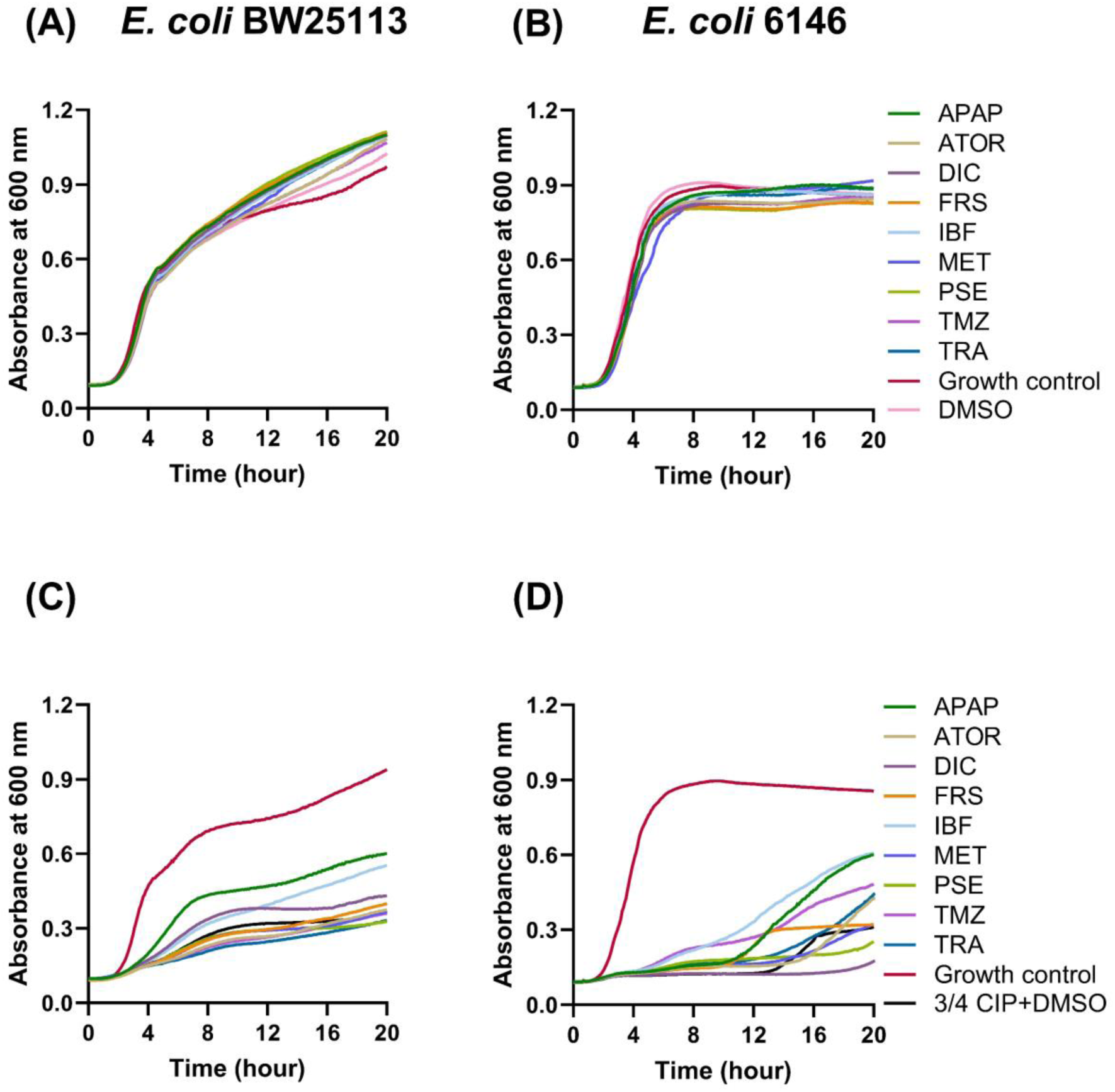
Growth profiles of *E. coli* BW25113 and 6146 in the presence of sub-inhibitory ciprofloxacin and NAMs. The effect of individual NAMs on growth of *E. coli* BW25113 (A) and *E. coli* 6146 (B) in the absence of ciprofloxacin and on the growth of *E. coli* BW25113 (C) and *E. coli* 6146 (D) in the presence of ¾ MIC ciprofloxacin. Growth control represented cells in the absence of NAMs and ciprofloxacin. Acetaminophen (APAP), atorvastatin calcium (ATOR), ciprofloxacin (CIP), diclofenac (DIC), furosemide (FRS), ibuprofen (IBF), metformin (MET), pseudoephedrine (PSE), temazepam (TMZ), tramadol (TRA). Growth curves were performed four times on different days with different batches of cells. Representative traces are shown here.

### 3.2. NAM exposure induces mutations within E. coli

To investigate the potential of the NAMs assessed in this study to induce the development of AMR by mutation, *E. coli* BW25113 and 6146 cells were exposed to NAMs at the projected gut concentrations (Maier et al., 2018) and ciprofloxacin at 1 x the MIC (0.015 µg/mL) and cultured for 48 hours (**Figure 2A**). All assays were carried out on plates supplemented with 0.015 µg/ml ciprofloxacin, as analysis of growth at higher ciprofloxacin concentrations revealed no growth.

**Figure 2.**
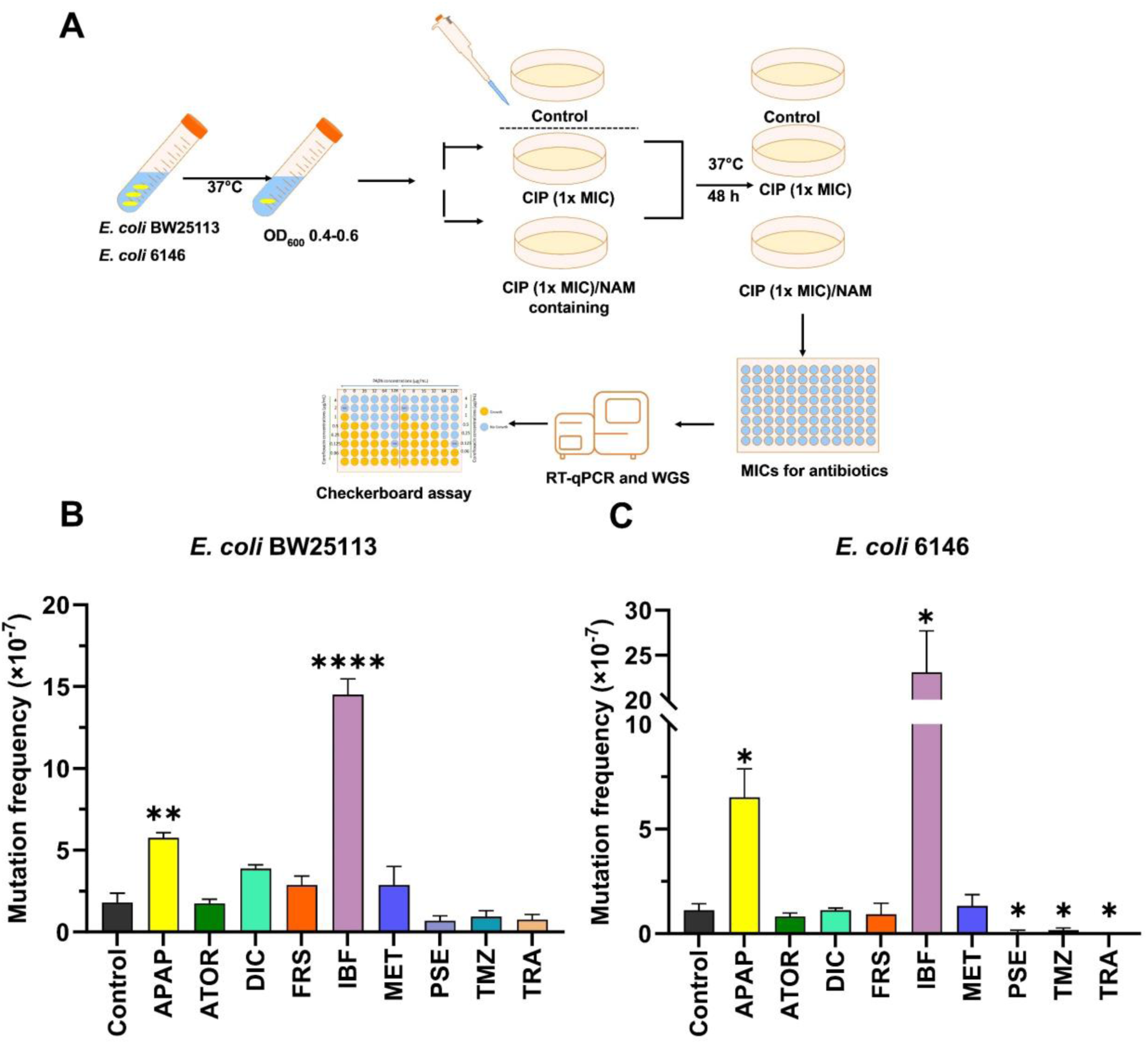
Mutation frequencies following exposure to ciprofloxacin and NAMs. (A) Workflow used to determine the mutation frequencies of *E. coli* BW25113 and 6146 cells exposed to NAMs and ciprofloxacin. The mutation frequency of *E. coli* BW25113 (B) and *E. coli* 6146 (C) were determined upon the exposure to NAMs at 216 μg/mL for acetaminophen (APAP), 6.6 μg/mL for atorvastatin (ATOR), 15.4 μg/mL for diclofenac (DIC), 13.2 μg/mL for furosemide (FRS), 99.8 μg/mL for ibuprofen (IBF), 194.6 μg/mL for metformin (MET), 40 μg/mL for pseudoephedrine (PSE), 3.3 μg/mL for temazepam (TMZ), and 33.3 μg/mL for tramadol (TRA) and ciprofloxacin (CIP) (0.015 μg/mL). Differences between mutants treated with NAMs and the Control (ciprofloxacin treated only) were analysed by independent-sample t-tests and corrected by Bonferroni correction method, *P < 0.05, **P < 0.01, ***P < 0.001, and ****P < 0.0001. Frequencies were averaged from four experiments done on different days with different batches of cells.

Higher mutation frequencies were observed for *E. coli* BW25113, with (1.45 ± 0.19) × 10^-6^ and (5.75 ± 0.64) × 10^-7^ cells exposed to ibuprofen (P < 0.0001) and acetaminophen (p<0.01), respectively, compared to exposure to ciprofloxacin alone (**Figure 2**). Similarly, a significant increase in mutation frequency was observed for *E. coli* 6146 following the exposure to acetaminophen (6.50 ± 2.73) × 10^-7^ and ibuprofen (2.30 ± 0.92) × 10^-6^. Remarkably, the mutation frequencies of the pseudoephedrine, temazepam, and tramadol showed a significant decrease for *E. coli* 6146 relative to exposure to ciprofloxacin only. The mutation frequencies of the remaining NAMs assessed in this study, were not significantly different from that of the ciprofloxacin only treatment, though the mutation frequencies for *E. coli* BW25113 was higher than for the clinical strain when cells were exposed to ciprofloxacin in combination with diclofenac, atorvastatin, furosemide, and metformin (**Figure 2**).

### 3.3. NAM exposure leads to elevated ciprofloxacin resistance in select isolates

To investigate levels of acquired ciprofloxacin resistance, a representative number (√n+1) of *E. coli* BW25113 and 6146 derived mutants were selected from each ciprofloxacin/NAM combination plate and the MIC of ciprofloxacin was determined (**Figure 3)**. As expected, *E. coli* BW25113 cells exposed to ciprofloxacin only developed resistance resulting in increased MICs of ciprofloxacin (2 to 8-fold increase in ciprofloxacin MIC as compared to WT *E. coli*). A comparable rise in ciprofloxacin resistance was also observed for NAM/ciprofloxacin derived mutants, with most *E. coli* BW25113-derived mutants displaying a 2 to 4-fold increase in MICs (**Figure 3A**). However, two mutants, one exposed to atorvastatin/ciprofloxacin (Ator_M1) and one exposed to ibuprofen/ciprofloxacin (Ibf_M1) displayed higher levels of ciprofloxacin resistance with a MIC of 0.125 µg/mL (8-fold increase). Two other BW25113 *E. coli* derived mutants, acetaminophen/ciprofloxacin Apap_M1 and Apap_M2 displayed a 16 and 4-fold increase in ciprofloxacin resistance respectively. Interestingly, the ibuprofen/ciprofloxacin and acetaminophen/ciprofloxacin not only displayed a high increase in ciprofloxacin MIC, but the mutation frequencies for these NAM/ciprofloxacin combinations were the only ones that were significantly higher than the mutation frequency of ciprofloxacin alone. Therefore, these mutants were selected for further assessment and their resistance profiles against an extended number of antibiotics determined.

**Figure 3.**
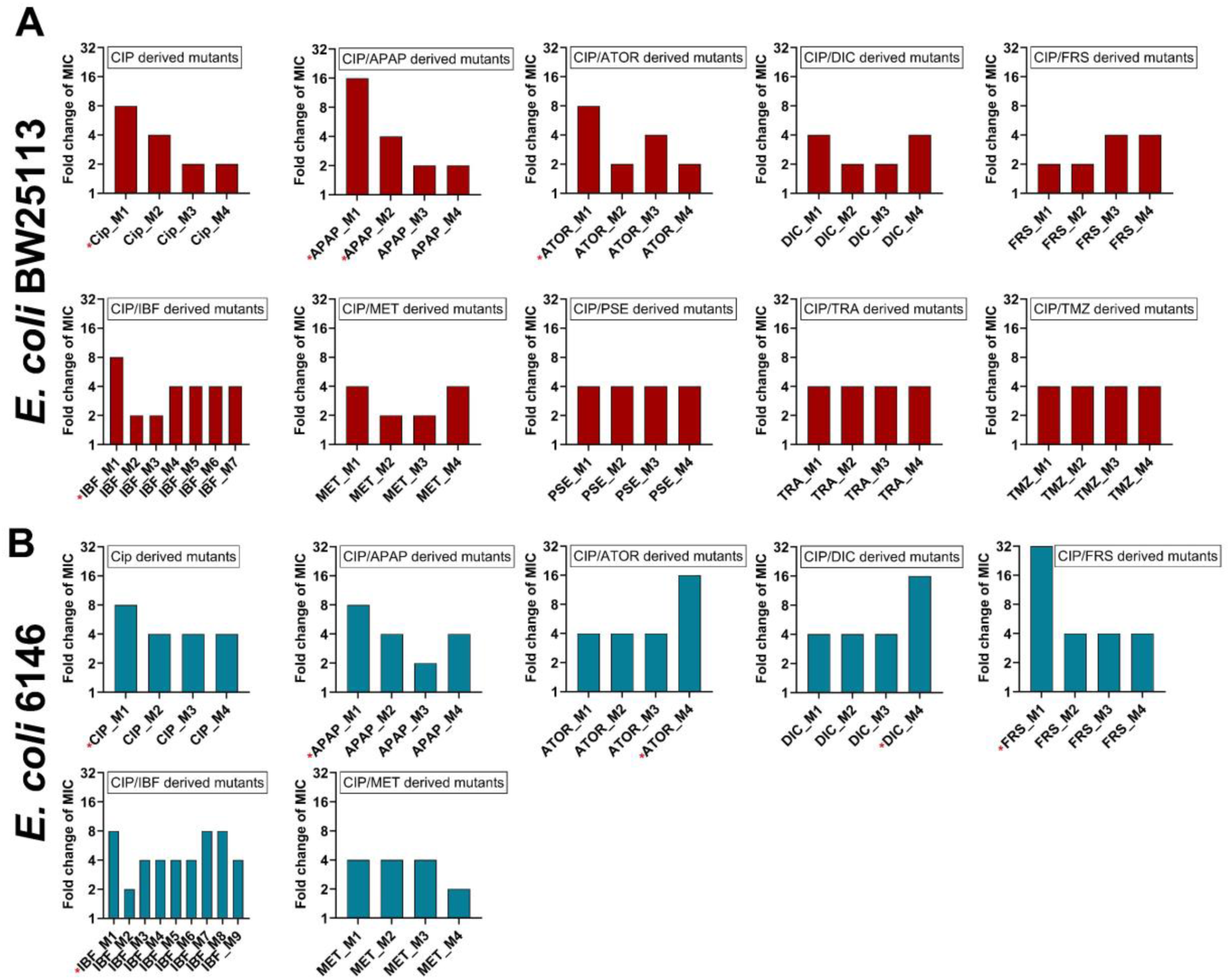
Fold change of MIC values for ciprofloxacin against NAM/ciprofloxacin mutants compared to WT. A representative number (√n+1) of *E. coli* derived mutants from select NAM/ciprofloxacin plates were picked and the MIC of ciprofloxacin was determined for *E coli* BW25112 (A) and *E coli* 6146 (B). * Indicates mutants selected for further analysis.

Evaluation of the *E. coli* 6146 derived mutants and an examination of the MICs against ciprofloxacin revealed a similar trend, with ciprofloxacin only exposed mutants showing a 4 to 8-fold increase in ciprofloxacin MIC as compared to the wild type. Mutants exposed to pseudoephedrine, temazepam, and tramadol/ciprofloxacin were not further investigated due to the limited number of colonies on plates. Notably, higher ciprofloxacin MIC values were observed for the *E. coli* 6146-derived mutants when compared to BW25113-derived mutants. The ciprofloxacin MIC for one mutant exposed to furosemide/ciprofloxacin (Frs_M1) increased to 0.5 µg/mL (32-fold increase in MIC), hence this mutant was selected for further assessment. Also selected for further analysis were mutants Ator_M4 (atorvastatin/ciprofloxacin exposed) and Dic_M4 (diclofenac/ciprofloxacin exposed) as these also displayed higher levels of ciprofloxacin resistance (MICs of 0.25 µg/mL) displaying 16-fold increase compared to WT *E. coli* 6146, as well and one acetaminophen/ciprofloxacin mutant (6146_Apap_M2) with a MIC of 0.125 µg/mL, showing an 8-fold increase ciprofloxacin resistance. Finally, as a larger mutation frequency was observed for cells following ibuprofen/ciprofloxacin exposure, a representative mutant, 6146_Ibf_M1, which displayed a MIC of 0.06 µg/mL (4-fold increase in MIC) was also assessed further.

### 3.4. NAMs exposure increases resistance to multiple antibiotics

Evaluation of selected ciprofloxacin resistant *E. coli* BW25113 and 6146 derived mutants against select antibiotics (**Table S1**), revealed elevated levofloxacin resistance for a large number of mutants. This is expected as the same resistance mechanisms are utilised by *E. coli* against these quinolones (Morgan-Linnell et al., 2009). However, the fold increase in MIC values for ciprofloxacin or levofloxacin differed for the mutants assessed here, suggesting a slight variation in the mechanisms of resistance against these two compounds (Fu et al., 2013). Of the compounds assessed here, in addition to increased ciprofloxacin and levofloxacin MICs, all BW25113-derived mutants except for M1 and Ator_M1 displayed a 4-fold increase in the MIC for minocycline. A 4-fold increase in the MIC for ceftazidime was also observed for Apap_M2 and Ator_M1.

Evaluation of resistance of the clinical *E. coli* 6146-derived mutants revealed higher MIC values of ciprofloxacin compared to the mutants derived from the *E. coli* BW25113 strain. For example, mutant Frs_M1 saw an increase of 32-fold for ciprofloxacin and 16-fold for levofloxacin. A 16-fold increase in MIC for ciprofloxacin was also observed for mutants Dic_M4 and Ator_M4. These mutants also showed elevated levofloxacin resistance with an 8-fold increase in their MICs against this compound. In addition, elevated resistance against cefepime was observed in the M1 mutant, Apap_M1, Dic_M1, and Ator_M1, with a 4-fold increase in the MIC of cefepime. Comparatively, while mutants derived from *E. coli* 6146 displayed higher levels of resistance against ciprofloxacin, levofloxacin, and cefepime than the BW25113 strain, resistance to other antibiotics such as ceftazidime and minocycline were not observed. Although Frs_M1 did not show high mutation frequency, the 32-fold increase in MIC of ciprofloxacin qualified it for further assessment.

### 3.5. The combination of two NAMs increases fitness in the presence of sub-MIC concentration of ciprofloxacin

To further explore the effects of a combination of medications, mutation frequencies were evaluated in the presence of two NAMs and ciprofloxacin. As exposure of *E. coli* BW25113 and 6146 to gut concentrations of ibuprofen and acetaminophen individually significantly enhanced mutation frequencies, and as mutants derived from the exposure to these compounds in the presence of ciprofloxacin displayed elevated MIC values for ciprofloxacin and other classes of antibiotics, these two NAMs were chosen for further analysis, and their ability to induce mutations were assessed in the presence of an additional NAM.

Bacterial growth was assessed in the presence of ibuprofen- and acetaminophen-based NAM combinations (**Figure 4**). The growth of *E. coli* BW25113 and *E. coli* 6146 cells was not significantly affected when these two strains were exposed to two NAMs in the absence of ciprofloxacin. For *E. coli* BW25113, acetaminophen-based combinations, with the exceptions of APAP+TMZ and APAP+MET, increased the growth rate, whereas most ibuprofen-based combinations decreased the growth rate. For *E. coli* 6146, the addition of NAM combinations, especially for the ibuprofen-based NAM combinations (except for IBF+FRS), resulted in an increase in growth rate, maximum cell density and a reduction in the log phase delay compared to NAM-free control (3/4 CIP + DMSO). The results indicate that these NAM combinations could enhance the fitness and adaptability of these two strains.

**Figure 4.**
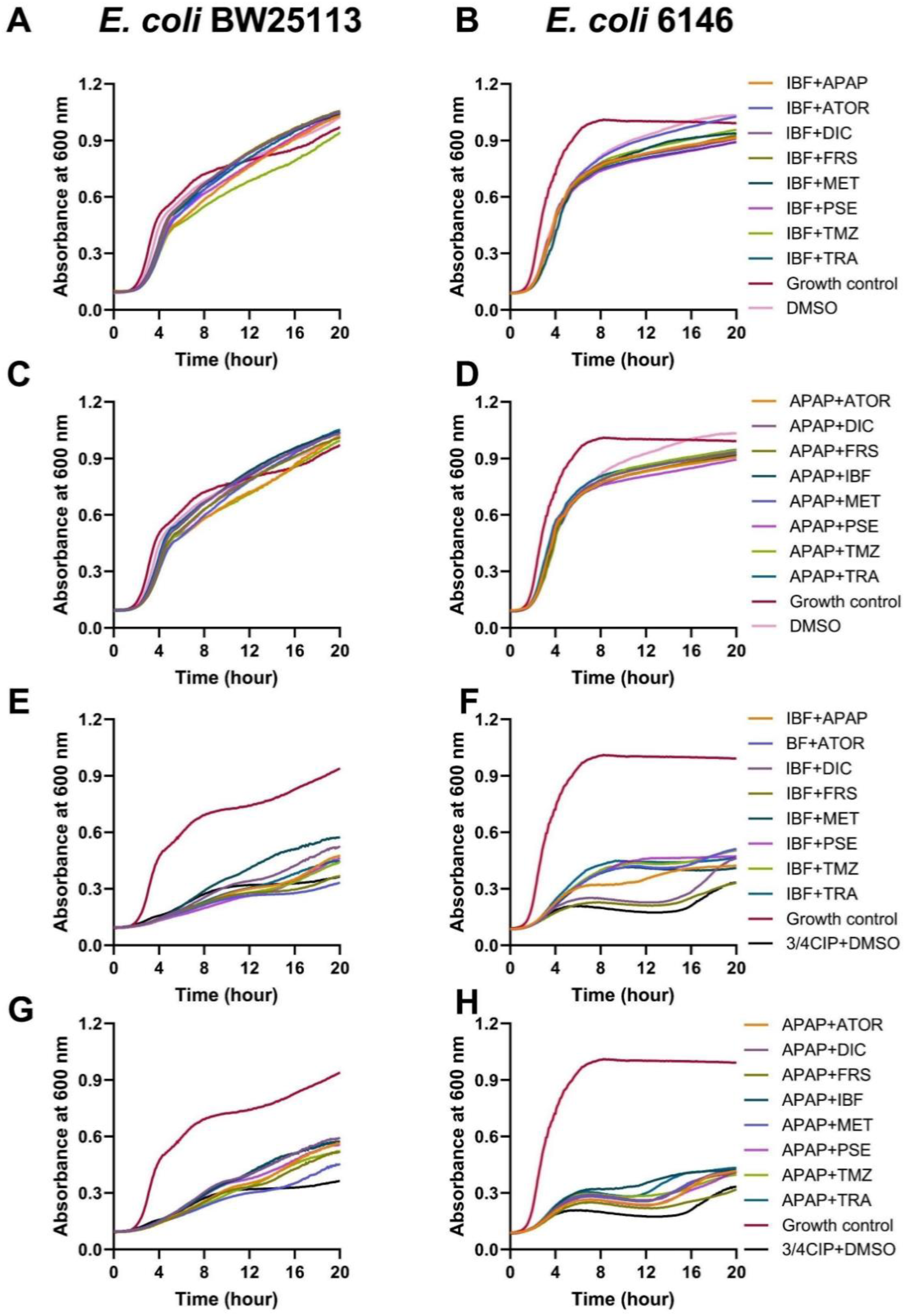
Growth profiles of *E. coli* BW25113 and *E. coli* 6146 in the presence of a sub-inhibitory inhibitory concentration of ciprofloxacin and 2 NAMs. The effect of 2 NAMs on growth of *E. coli* BW25113 (**A** and **C**) and *E. coli* 6146 (**B** and **D**). The effect of 2 NAMS on the growth of *E. coli* BW25113 (**E** and **G**) and *E. coli* 6146 (**F** and **H**) in the presence of ¾ MIC ciprofloxacin. Growth control represented cells in the absence of NAMs and ciprofloxacin. Acetaminophen (APAP), atorvastatin calcium (ATOR), ciprofloxacin (CIP), diclofenac (DIC), furosemide (FRS), ibuprofen (IBF), metformin (MET), pseudoephedrine (PSE), temazepam (TMZ), tramadol (TRA). Growth curves were performed four times on different days with different batches of cells. Representative traces are shown here.

### 3.6. The combination of two NAMs enhances the mutation frequency in the presence of ciprofloxacin

Assessment of mutation frequency following the exposure of cells to 2 NAMs and ciprofloxacin, revealed that the presence of acetaminophen combined with ibuprofen significantly promoted the number of *E coli* BW25113 (P < 0.05) and 6146 (P < 0.01) mutants compared to cells exposed to ciprofloxacin only **(Figure 5)**. The mutation frequency for *E. coli* exposed to acetaminophen and ibuprofen (1.40 ± 0.56 × 10^-6^ for BW25113 and 1.57 ± 0.51 × 10^-6^ for 6146) was similar and even lower to that of cells exposed to ibuprofen alone (1.45 ± 0.19 × 10^-6^ for BW25113 and 2.30 ± 0.92 × 10^-6^ for 6146). Notably, a high mutation frequency (P < 0.05) was also observed for BW25113 cells following the exposure to ibuprofen and temazepam. As for *E. coli* 6146, the exposure to atorvastatin, diclofenac (P < 0.01), frusemide, and metformin (P < 0.05) resulted in the significantly increased mutation frequency compared to the cells exposed to ciprofloxacin only treatment. In contrast, the remain NAMs with ibuprofen showed decreased mutation frequency, relative to that of cells exposed to ibuprofen.

**Figure 5.**
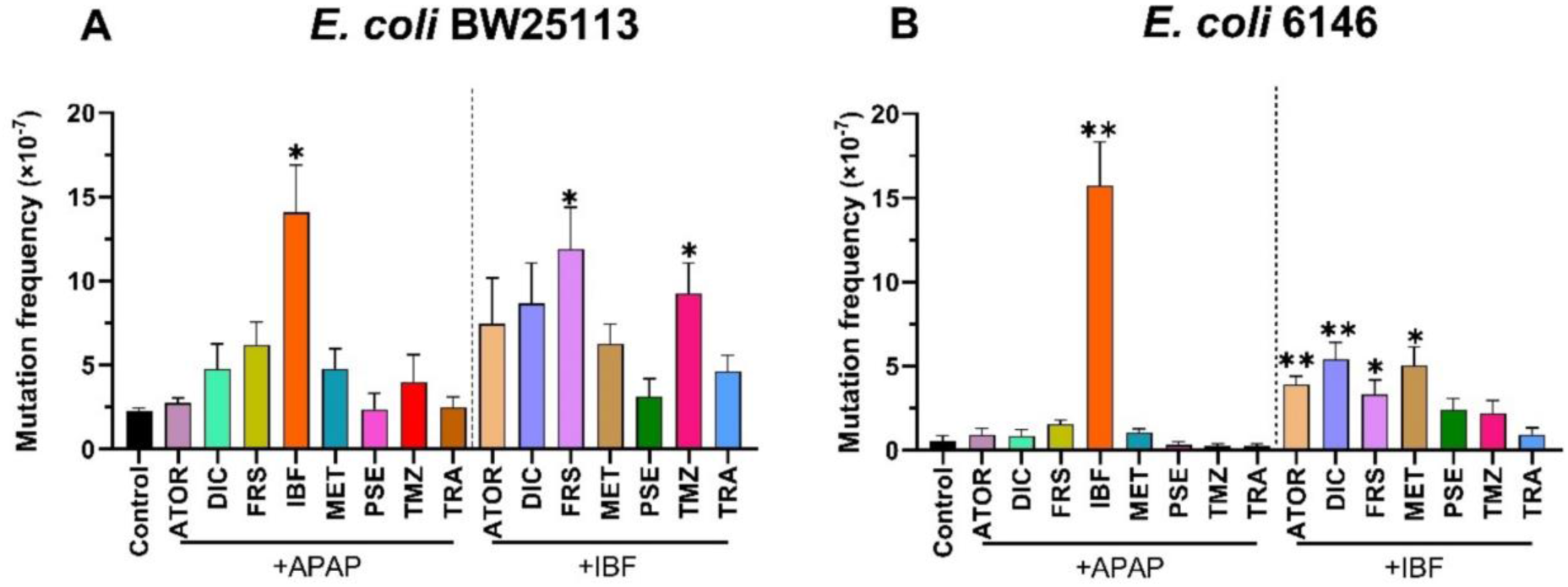
Mutation frequencies of *E. coli* BW25113 and *E. coli* 6146 following exposure to ciprofloxacin and NAM combinations. The mutation frequency following the exposure to two NAMs are indicated for **A.** *E. coli* BW25113 and **B.** *E. coli* 6146. Significant differences between mutants treated with additional NAMs and the control (ciprofloxacin treated only) were analysed by independent-sample t-tests and corrected by Bonferroni correction method, *P < 0.05, **P < 0.01, and ****P < 0.0001. Acetaminophen (APAP), atorvastatin calcium (ATOR), ciprofloxacin (CIP), diclofenac (DIC), furosemide (FRS), ibuprofen (IBF), metformin (MET), pseudoephedrine (PSE), temazepam (TMZ), tramadol (TRA).

### 3.7. Exposure to two NAMs leads to elevated ciprofloxacin resistance in select isolates

A representative number (√n+1) of *E. coli* BW25113 and 6146 derived mutants were randomly picked from ciprofloxacin/NAMs combination plates and the MIC values for a panel of antibiotics were determined (**Figure 6)**. As expected, most mutants exposed to two NAMs/ciprofloxacin displayed higher levels of ciprofloxacin resistance, with a 2 to 4-fold increase in MICs compared to mutants exposed to ciprofloxacin and one NAM. In *E. coli* BW25113 derived mutants, two mutants, Apap/Ibf_M1 and Ibf/Met_M1, showed a 32-fold increase in ciprofloxacin resistance, with a MIC of 0.5 µg/mL. Notably, the mutant Ibf/Dic exhibited the highest levels of ciprofloxacin resistance, with an MICs of 1 µg/mL, representing a 64-fold increase compared to the MIC value of the WT.

**Figure 6.**
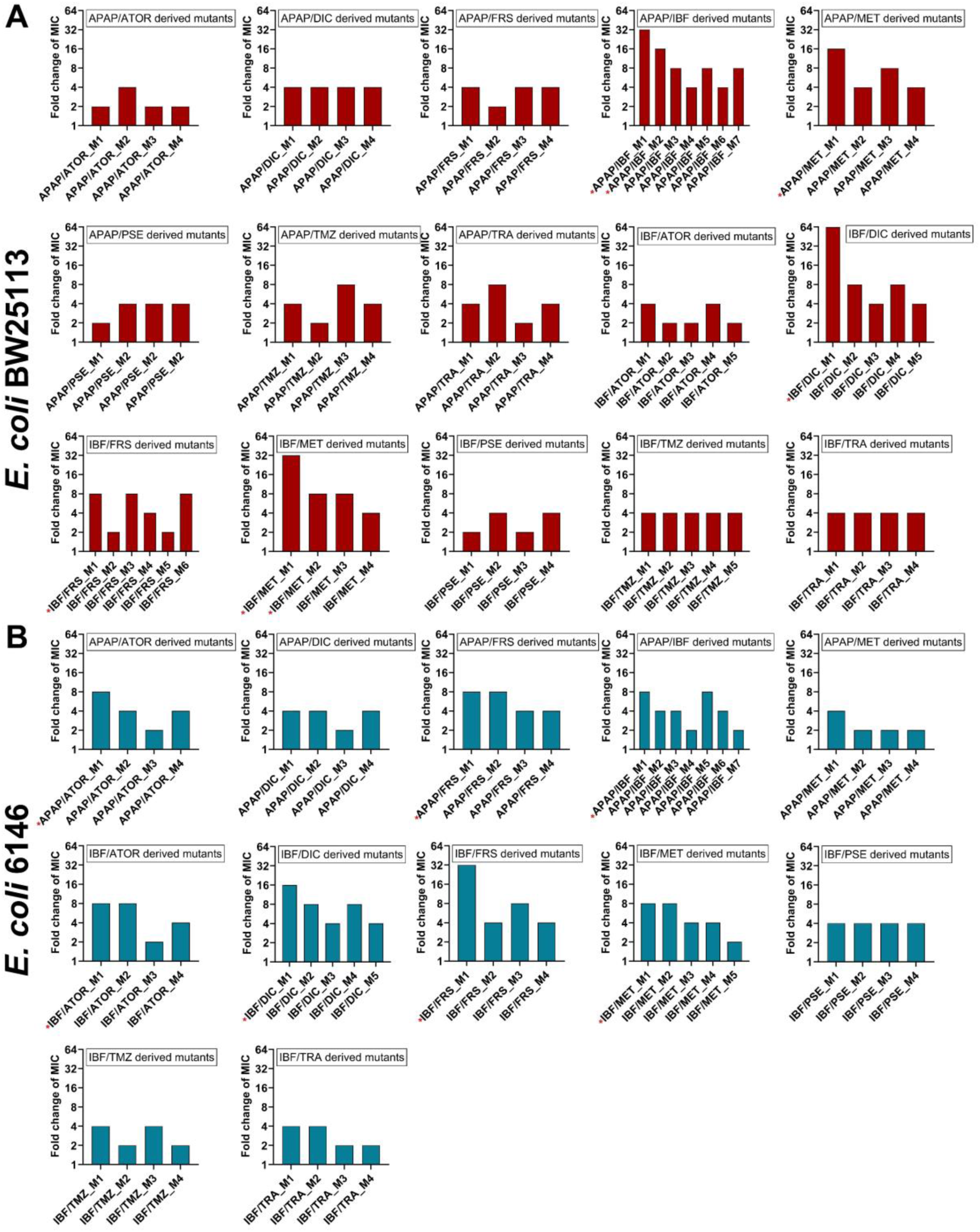
Fold change of MIC values for ciprofloxacin against two NAMs/ciprofloxacin mutants compared to WT. A representative number (√n+1) of *E. coli* derived mutants from select 2 NAMs/ciprofloxacin plates were picked and the MIC of ciprofloxacin was determined for *E coli* BW25112 (A) and *E coli* 6146 (B) derived mutants. Asterisks indicate mutants selected for further analysis. Acetaminophen (APAP), atorvastatin calcium (ATOR), ciprofloxacin (CIP), diclofenac (DIC), furosemide (FRS), ibuprofen (IBF), metformin (MET), pseudoephedrine (PSE), temazepam (TMZ), tramadol (TRA).

Concerning *E. coli* 6146 derived mutants, two mutants, Ibf/Frs_M1and Ibf/Dic_M1, exhibited higher ciprofloxacin resistance, with MICs of 0.5 µg/mL and 0.25 µg/mL, showing a 32- and 16-fold increase in ciprofloxacin MIC values respectively. The following mutants were selected for further assessment, Apap/Ibf_M1, Apap/Frs_M1, Apap/Ator_M1, and Ibf/Met_M1 from *E. coli* 6146 derived mutants, as MICs of 0.125 µg/mL were observed.

Mutants with at least 8-fold increased ciprofloxacin resistance representing each group of resistant isolates were selected and their resistance profiles against ten antibiotics were determined (**Table S2**). The mutants derived from exposure to two NAMs exhibited a higher level of ciprofloxacin resistance than those derived from the exposure to a single NAM, with such mutants as Ibf/Dic_M1 whose ciprofloxacin resistance increased up to 64-fold and mutants Apap/Ibf_M1 and Ibf/Met_M1 showing a 32-fold increase in ciprofloxacin MIC relative to wild type *E. coli* BW25113. Similarly, the MIC values for levofloxacin, an antibiotic from the same class as ciprofloxacin, also increased by 4 to 16-fold for the NAM derived mutants. In comparison, increase in resistance against antibiotics from other classes was lower, with mutants Apap/Met_M1, Ibf/Frs_M1 and Ibf/Met_M2 displaying an increase in MIC of 2 to 4-fold for minocycline, cefepime, ceftazidime and amoxicillin.

For *E. coli* 6146, mutants Ibf/Frs_M1 and Ibf/Dic_M1 manifested 32-fold and 16-fold increases in ciprofloxacin MIC values. These mutants also showed increased levofloxacin resistance, with a 16 and 8-fold increase in MIC, respectively. Finally, 4-fold increased cefepime resistance was observed in mutants Ibf/Apap_M1 and Apap/Ator_M1, with the latter also displaying increase minocycline resistance (4-fold increase in MICs).

### 3.8. Genomic analysis reveals multiple mutations in mutants exposed to NAMs and ciprofloxacin

To identify mutations which gave rise to ciprofloxacin resistance, mutants were selected for whole genome sequencing. The resistance determinants assessed were primarily focused on those involved in fluoroquinolone resistance, such as GyrA/B, ParC/E (Arzanlou et al., 2017; Redgrave et al., 2014) as well as on RND efflux pumps (AcrAB-TolC) (Olga Lomovskaya & Bostian, 2006; Venter et al., 2015), and their regulators (local regulator AcrR and global regulator MarR) (Piddock, 2006), as these also play an important role in such resistance (Amsalu et al., 2020). Finally, mutations within stress response proteins such as SoxS and SoxR were also assessed as these have previously been identified to play a role in the overexpression of efflux pumps (Koutsolioutsou et al., 2005).

Several mutations were identified in the mutants assessed, with mutations and deletions observed in MarR, AcrR and GyrA **(Table 2).** For the BW25113 derived mutant M1, a unique substitution mutation was detected in SoxR (R20L). Previously described GyrA mutations, S83L, D87G and D87Y, which are reportedly associated with a strong influence on quinolone resistance (Bagel et al., 1999; Fu et al., 2013) were observed in five mutants. Of these, one mutant was derived from *E. coli* 6146 cells exposed to frusemide and four from cells exposed to ibuprofen in addition to one other NAM and ciprofloxacin. All mutations related to efflux pumps mainly focus on their regulators, AcrR and MarR. Among these, a substitution mutation of AcrR (T5N) was identified in three mutants: Ibf_Frs and Apap_Met_M1 from BW25113, and Ibf/Apap_M1from 6146. Meanwhile, unique deletions in the MarR regulator occurred among these mutants. Interestingly, many mutants exposed to two NAMs exhibited more than one mutation *i.e*. an increase of mutation rate as well as mutation frequency was observed.

**Table 2.** Mutations, deletions and truncations of select proteins in *E. coli* BW25113 and *E. coli* 6146 derived mutants following the exposure to one and two NAMs and 1x MIC of ciprofloxacin.

|  | Strains | Protein | Mutation(s)* | Fold increase in Cip MIC |
| --- | --- | --- | --- | --- |
| BW25113 derived | BW_M1 | SoxR | R20L | 8 |
|  | BW_Ibf_M1 | ND | ND | 8 |
|  | BW_Apap_M1 | MarR | E136D, deletion – Y137 | 16 |
|  | BW_Apap_M2 | MarR | 1 base pair deletion | 4 |
|  | BW_Ator_M1 | MarR | deletions (L75, D76) | 8 |
|  | BW_Ibf/Met_M1 | GyrA | D87Y | 32 |
|  | BW_Ibf/Dic_M1 | GyrA | S83L | 64 |
|  | BW_Ibf/Frs_M1 | MarR | L78R | 8 |
|  |  | AcrR | T5N |  |
|  | BW_Apap/Ibf_M1 | GyrA | S83 deletion | 32 |
|  |  | MarR | V84E |  |
|  | BW_Apap/Met_M1 | MarR | 13 base pair deletion | 16 |
|  |  | AcrR | T5N |  |
| 6146 derived | 6146_M1 | MarR | 180 base pair deletion | 8 |
|  | 6146_Ibf_M1 | GyrA | 3 base pair deletion (S83 deletion) | 8 |
|  | 6146_Frs_M1 | GyrA | S83L | 32 |
|  | 6146_Ibf/Frs_M1 | GyrA | D87G | 32 |
|  |  | MarR | 13 base pair deletion |  |
|  | 6146_Ibf/Apap_M1 | MarR | 2 base pair insertion | 8 |
|  |  | AcrR | T5N |  |

ND: No mutations determined. Cip – ciprofloxacin, MIC – Minimum Inhibitory Concentration

### 3.6. Assessment of the role of efflux pumps in acquired resistance

Given that many mutations found in the *E. coli* derived mutants were related to efflux pump regulators, the potential role of RND efflux pumps in the observed ciprofloxacin resistance was investigated. Checkerboard assays were carried out in the presence of PAβN, an efflux pump inhibitor (Lomovskaya et al., 2001; Venter, 2019). In total, 15 mutants were assessed to determine the role of RND efflux pumps in resistance, with results revealing that the addition of PAβN partially reversed resistance (2-fold reduction) in four of the *E. coli* BW25113 derived mutants, with a further 5 mutants seeing a 4-fold reduction in ciprofloxacin MIC and one mutant, Ibf/Frs_M1, displaying an 8-fold reduction in ciprofloxacin resistance (**Figure 7A**). All BW25113 derived mutants displaying a 4 to 8-fold reduction in ciprofloxacin resistance were found to carry mutations in MarR or AcrR or both. Finally, the addition of PAβN to two BW25113 derived mutants, Apap_M2 and Ibf/Frs_M1, abolished ciprofloxacin resistance, indicating that the resistance observed is mediated by efflux through the RND-type efflux pumps.

**Figure 7.**
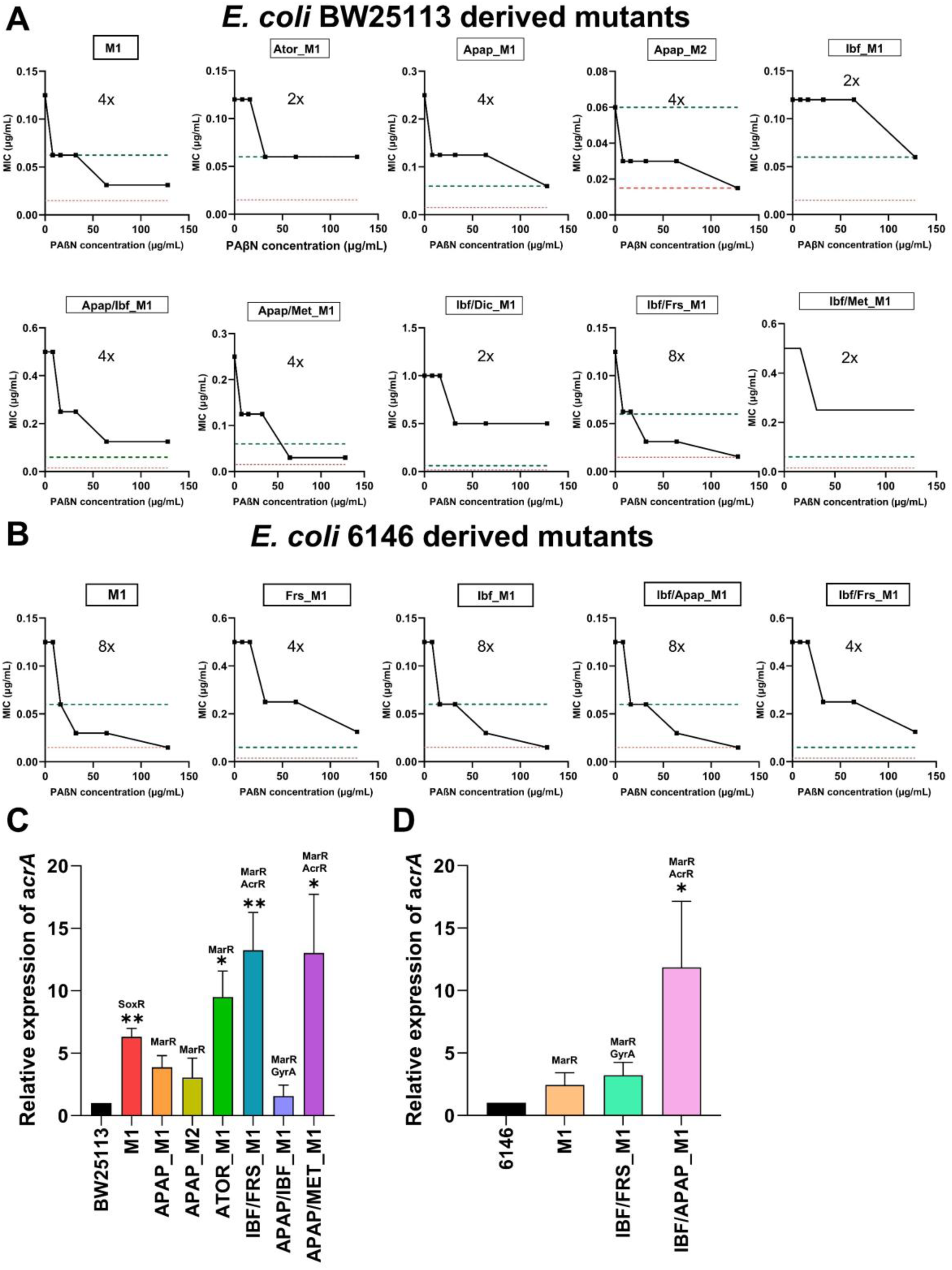
Evaluation of efflux pump-mediated antimicrobial resistance. Reversal of resistance in the presence of the efflux pump inhibitor PAβN for *E. coli* BW25113 (A) and *E. coli* 6146 (B) derived mutant isolates. The black lines correspond to the MICs in the presence of PAβN, and red and green dashed lines represent the MIC of wild type and average MIC of mutants exposed to ciprofloxacin only respectively. The expression levels of the AcrAB-TolC efflux pump were assesses by RT-qPCR of the *acrA* gene in *E. coli* BW25113 (C) and *E coli* 6146 (D) derived mutants. The results presented are the mean ± SEM of three experiments with different batches of cells and RNA preparations. Significant differences between mutants and the WT strain were analysed by independent-sample t-tests and corrected by Bonferroni correction method, *P < 0.05, **P < 0.01, ***P < 0.001, and ****P < 0.0001. Acetaminophen (APAP), atorvastatin calcium (ATOR), ciprofloxacin (CIP), diclofenac (DIC), furosemide (FRS), ibuprofen (IBF), metformin (MET).

The addition of PAβN to *E. coli* 6146 derived mutants resulted in greater reductions in resistance, with mutants Frs_M1 and Ibf/Frs_M1 displaying 4-fold reductions in ciprofloxacin MIC, and an 8-fold reduction in resistance seen in the remaining three mutants **(Figure 7B).** The addition of PAβN to three *E. coli* 6146 derived mutants, also saw a complete reversal in ciprofloxacin resistance. Of these, Ibf_M1 was found to harbor a GyrA mutation (ΔS83), and no additional efflux pump associated mutations. As such the reversal of resistance may be due to the transient upregulation of efflux pumps (Arzanlou et al., 2017; Hernando-Amado et al., 2016), which was subsequently inhibited by the addition of PAβN.

### 3.9. RT-qPCR revealed AcrAB-TolC efflux pump over-expression

As numerous MarR and AcrR mutations were observed in many of the NAM derived *E. coli* mutants and reduction in MIC values were observed in the presence of the efflux pump inhibitor PAβN, RT-qPCR was carried out to assess the upregulation of the AcrAB-TolC efflux pump. A total of ten NAM derived *E. coli* mutants, seven *E. coli* BW25113 and three *E. coli* 6146 were selected for analysis **(Figure 7).** A significant increase in the expression of the AcrAB efflux pump was observed in all the mutants showing a fold increase ranging from 2.44 to 13.02 compared to their wild type strains, with the exception of the BW25113-derived mutant Apap/Ibf_M1. The BW25113-derived mutant Apap/Ibf_M1 showed a 4-fold reduction in ciprofloxacin MIC following the addition of PAβN and harboured a mutation in MarR (V84E), but the expression levels of AcrA only increased by 1.56-fold. Notably, the most substantial increase in AcrAB expression was observed in BW25113 derived mutants Ibf/Frs_M1 (P < 0.001), Apap/Met_M1 (P < 0.001), and 6146 derived mutant Ibf/Apap_M1 (P < 0.01), exhibiting 13.24-, 13.02- and 11.85-fold upregulation, respectively. Interestingly, these mutants harboured MarR as well as the AcrR (T5N) mutations. In addition, the *acrA* gene encoding efflux pump was significantly upregulated in mutant Aror_M1 containing a 2bp deletion in MarR (P < 0.05).

## Discussion

Since their discovery, antibiotics have remained essential in the treatment of infectious diseases worldwide. However, their overuse and misuse have been a major driving force behind decreased antibiotic susceptibility and increased antibiotic resistance (Imperial & Ibana, 2016). This phenomenon is especially prevalent in RACFs, where inappropriate use of antibiotics are prevalent and where the prevalence of resistant bacteria continues to grow (Blaikie et al., 2024; Lim et al., 2014; Sapula et al., 2023; Sapula et al., 2024; Stuart et al., 2012). Compounding this is the fact that in addition to antibiotics, residents are also prescribed numerous NAMs which can also contribute to the development of resistance within bacteria. These drugs include the non-steroidal anti-inflammatory drugs (NSAIDs) ibuprofen and diclofenac, opioid medication tramadol, and the non-opioid analgesic acetaminophen (also known as paracetamol) which are used for pain relief (Nowak et al., 2019). Pseudoephedrine, a sympathomimetic drug used for relieving nasal or sinus congestion, temazepam used to manage insomnia, metformin, a glucose lowering medication commonly used for the management of type 2 diabetes (Keitaanpaa et al., 2018; Poudel et al., 2016), furosemide, a loop diuretic used to manage oedema, and atorvastatin, a lipid lowering medication (i.e., 3-hydroxy-3-methylglutaryl-coenzyme A (HMG-CoA) reductase inhibitor) (Okafor et al., 2024), are also frequently used.

Previous studies have shown that antibiotics at sub-inhibitory concentrations could promote the development and spread of antibiotic resistance (Gullberg et al., 2011; Sanz-García et al., 2022). However, little is known about the contribution of NAMs to the emergence and spread of AMR. To address this, we screened a panel of nine NAMs against *E. coli* to assess their effect on the mutation frequency in *E. coli*. Collectively, the addition of certain NAMs like ibuprofen, acetaminophen, and diclofenac led to enhanced mutation frequencies, relative to cells exposed to ciprofloxacin only. In addition, ibuprofen, and acetaminophen, amongst others were found to protect the cells from ciprofloxacin-induced injury during the growth phase, as evidenced by an increased growth rate and maximum cell density. Generally, the long-term consumption of medications among older persons leads to combined exposure to antibiotics like ciprofloxacin and NAMs in the range of mg/L in the gut microbiota (Maier et al., 2018; Shi et al., 2022). These findings highlight the potential for *E. coli* to develop mutations leading to AMR following exposure to NAMs and ciprofloxacin in RACFs where medications are widely used.

Overall, our results demonstrated that two commonly consumed pain relief medications (ibuprofen and acetaminophen), can significantly enhance antibiotic resistance via mutation at gut-relevant concentrations in *E. coli* strain BW25113 and *E. coli* 6146. This was evident in the increased number of mutants and the enhancement of maximum cell density. Other NAMs such as diclofenac and furosemide, also increased mutation frequency when compared to the non-NAM control (only ciprofloxacin). It has been reported that frequent exposure to antidepressants such as fluoxetine, sertraline, and duloxetine could induce AMR through the mutations (Wang et al., 2023). However, the other NAMs in that study (temazepam, tramadol, and pseudoephedrine) had minimal impact on mutation frequency for *E. coli* BW25113 and even exhibited negative effects for *E. coli* 6146. Considering polypharmacy is common among the elderly in RACFs, the effects caused by ibuprofen and acetaminophen also warrants attention. Interestingly, we found that the combined exposure to ibuprofen and acetaminophen induced similar mutation frequency to that of ibuprofen alone, rather than resulting in a cumulative increase in mutation frequency.

In addition, *E. coli* BW25113 and 6146 derived mutants exhibited reduced susceptibility to multiple antibiotics (≥4-fold), particularly showing increased ciprofloxacin resistance, with MIC values rising to 32-fold for this agent. Among these antibiotics, β-lactam antibiotics (amoxicillin, ceftazidime, meropenem), as well as ciprofloxacin, levofloxacin, and minocycline are substrates of the AcrAB-TolC efflux pump, which can extrude multiple classes of antibiotics (Du et al., 2014; Kobylka et al., 2020). Therefore, increased resistance observed against these antibiotics suggests a potential link to this efflux pump. Furthermore, RT-qPCR analysis revealed the overexpression of AcrAB-TolC efflux pump, supporting a close relationship between the NAMs and efflux pump expression. Previous research has suggested that human-target pharmaceuticals may boost antibiotic resistance through the regulation of efflux pumps (Maier et al., 2018). In this study, although exposure to two NAMs did not induce higher mutation frequencies, mutants derived from two NAMs/ciprofloxacin combinations exhibited higher levels of resistance to multiple antibiotics, in particular to ciprofloxacin compared to those exposed to a single NAM/ciprofloxacin combination.

In this study, whole genome sequencing was conducted to elucidate the mechanisms responsible for multiple antibiotic resistance in mutants. The analysis revealed that the combination of NAMs with ciprofloxacin can induce nonsynonymous single nucleotide polymorphisms (SNPs) in regulator genes, leading to the efflux pump overexpression. In addition to mutations occurring in quinolone resistance determinant regions in topoisomerases such as *gyrA,* multiple mutations were also identified (deletions and substitution) in regulator genes, such as *marR*, *acrR,* and *soxR* which are regulators of the AcrAB-TolC system, highlighting the connection between antibiotics and efflux pumps (Bialek-Davenet et al., 2011; Sanz-García et al., 2022). MarR negatively regulates the transcription factor MarA, which in turn regulates the expression of multiple other proteins including those of the AcrAB-TolC efflux pump. Therefore, the mutations in MarR resulted in increased expression of the marRAB operon, leading to upregulation of AcrAB-TolC efflux pump and reduced antibiotic susceptibility (Praski Alzrigat et al., 2017). Such as mutant BW_APAP/IBF carrying *marR* V84E and *gyrA* Δ83 not only conferred high ciprofloxacin resistance but also had a small increase in MICs to ceftazidime and minocycline. This was consistent with a previous study that clinical isolates carrying *marR* V84E provided antibiotic resistance to different classes of antibiotics (Ching & Zaman, 2020). AcrR acts as a local repressor to prevent the overexpression of the AcrAB-TolC efflux pump. Strikingly, upon exposure to two NAMs/ciprofloxacin, an increase in the number of mutations was observed in certain *E. coli* isolates, with five mutants developing mutations in both MarR and AcrR or GyrA and MarR. Of these, the T5N mutation observed in AcrR occurred in three mutants which showed reduced susceptibility to different classes of antibiotics. Also, RT-qPCR analysis showed mutants carrying the T5N substitution resulted in the highest level of AcrAB-TolC efflux pump expression. A previous study postulated that the T5N mutation observed in AcrR may render this regulator inactive or potentially destabilize it (Gerken & Misra, 2004), with the increased expression of the AcrAB-TolC efflux pump observed in this study, supporting this theory. Some global regulators such as SoxS, are regulated by SoxR, whose activation by superoxides contributes to the expression of SoxS, which in turn upregulates the transcription of *acrAB* (Aly et al., 2015; Pérez et al., 2012). The R20L substitution observed in the BW_M1 mutant could account for the upregulation of AcrAB-TolC efflux pump, thereby leading to increased resistance to antibiotics.

Our findings based on phenotypic analysis revealed that only a few mutants fully reverse ciprofloxacin resistance to the original levels in the presence of PAβN, an efflux pump inhibitor. It has been shown that high levels of antibiotic resistance are not only attributed to increased expression of efflux pumps but may result from the accumulation of mutations in different targets (Amsalu et al., 2020; Moon et al., 2010; Piddock, 2006). In this study, a combination of *gyrA* mutations along with mutations in efflux pump regulators contributed to the observed fluoroquinolone resistance. It is also thought that the mechanism enabling *E. coli* to survive in the presence of antibiotics is linked to the repression of DNA mismatch repair gene *mutS*, which is caused by upregulation of efflux pumps. This repression leads to a higher mutation frequency rate and explains how efflux pump overexpression facilitates the development of AMR (Darby et al., 2023; El Meouche & Dunlop, 2018). Microbes increase the efflux of antimicrobials to lower the intracellular concentration of antimicrobials and provide fitness, allowing them to survive longer in the presence of low concentrations of antibiotics than expected. (Piddock, 2006; Praski Alzrigat et al., 2017). Additionally, mutations in clinical isolates tend to favour those that impose small cost fitness and conferred small reduction in antibiotic susceptibility. This growth advantage allows the strain to gradually accumulate mutations, eventually evolving into high-level resistance (Piddock, 2006; Praski Alzrigat et al., 2017).

Collectively, this study provides evidence that medication used in pain relief such as acetaminophen and ibuprofen, when combined with ciprofloxacin, not only significantly increase mutation frequency but also confer multiple antibiotic resistance. Other NAMs like diclofenac and furosemide also contribute to increased mutation frequency rates and high levels of fluoroquinolone resistance. Although the mutation frequency rate induced by two NAMs is not synergistic for mutation frequency increase, high-level ciprofloxacin resistance is observed among the mutants. The accumulation of mutations is worrisome as these have been observed in mutants exposed to two NAMs and ciprofloxacin; However, the elderly in RACFs are taking numerous medications daily (Chen et al., 2019) highlighting the need to consider medicine combinations when prescribing. This study also paves the way for investigating the effect of regimens comprising multiple medications (i.e., polypharmacy) on the development of AMR.

## CRediT authorship contribution statement

**Hanbiao Chen:** Writing – original draft, Writing – review & editing, Methodology, Investigation, Formal analysis, Validation, visualization, Data curation,

**Sylvia A. Sapula:** Conceptualization, Writing – original draft, Writing – review & editing, Methodology, Investigation, Formal analysis, Resources, supervision, Validation, Data curation,

**John Turnidge**: Writing – review & editing

**Henrietta Venter:** Conceptualization, Methodology, Writing – review & editing, Funding acquisition, Project administration, Resources, supervision

## Acknowledgments

We are grateful to the aged care provider and Mr Barry Lowe for sample collection and lists of commonly used drugs in the RACFs and to A/Prof Janet Slugget (UniSA) for advice on polypharmacy and the use of medications in RACFs in Australia. We also thank the Dr. Lex Leong (SA Pathology) for providing whole genome sequencing.

